# Chemical potential measurements constrain models of cholesterol-phosphatidylcholine interactions

**DOI:** 10.1101/2022.10.08.511420

**Authors:** Thomas R. Shaw, Kathleen Wisser, Taylor A. Shaffner, Anna D. Gaffney, Benjamin B. Machta, Sarah L. Veatch

## Abstract

Bilayer membranes composed of cholesterol and phospholipids exhibit diverse forms of non-ideal mixing. In particular, many previous studies document macroscopic liquid-liquid phase separation as well as nanometer-scale heterogeneity in membranes of phosphatidylcholine (PC) lipids and cholesterol. Here, we present experimental measurements of cholesterol chemical potential (μ_c_) in binary membranes containing dioleoyl PC (DOPC), 1-palmitoyl-2-oleoyl PC (POPC), or dipalmitoyl PC (DPPC), and in ternary membranes of DOPC and DPPC, adapting a calibrated experimental protocol developed to measure μ_c_ in cells (Ayuyan and Cohen, Biophys. J. 114:904-918). μ_c_ is the thermodynamic quantity that dictates the availability of cholesterol to bind other factors, and notably must be equal between coexisting phases of a phase-separated mixture. It is simply related to concentration under conditions of ideal mixing but is found to be far from ideal for the majority of lipid mixtures investigated. Here we perform experimental measurements of μ_c_, constraining thermodynamic models of membrane interactions. Our measurements are consistent with models involving cholesterol-phospholipid complexes, but only if complexes are more weakly bound than has been assumed in previous reports. Experimental measurements are also well described by regular solution theory and lattice models with pairwise interactions between components. We find that μ_c_ can vary by ~1.5 *k*_*B*_*T* at constant cholesterol mole-fraction implying a more than five-fold change in its availability for binding receptors and other reactions. These findings reinforce that μ_c_ depends on membrane composition overall, suggesting avenues for cells to alter the availability of cholesterol without varying cholesterol concentration.

**SIGNIFICANCE:** The chemical potential of cholesterol (μ_c_) reflects its availability to interact with other molecules. In a complex mixtures this chemical potential can vary dramatically even at fixed cholesterol concentration. In this report, we present measurements of μ_c_ in bilayer membranes composed of mixtures of cholesterol with one or two phospholipids. We find that μ_c_ in these mixtures depends strongly on the phospholipids that are present, with activity varying by a factor of more than five at fixed cholesterol concentration. This suggests that the availability of cholesterol in biological membranes could be tuned without altering cholesterol concentration directly, by adjusting the concentration of other lipid or protein components.

## INTRODUCTION

Phospholipid bilayer membranes containing cholesterol are complex fluids that exhibit non-ideal mixing of components that is detectable by a broad range of experimental methods. Non-ideal mixing can take the form of phase separation or meso-scale nano-domains detected by methods such as Fourier resonance energy transfer (FRET), electron spin resonance (ESR), or nuclear magnetic resonance (NMR), neutron scattering, or fluorescence microscopy (e.g. (1–8) reviewed in (9)). This non-ideality is a ubiquitous feature of these membranes and therefore it is expected to contribute to their chemical and material properties.

One fundamental biophysical property of components within membranes is their chemical potential (μ), which governs, among other things, how available these components are to bind to embedded proteins. The quantity exp{μ / *k*_*B*_*T*}is called the chemical activity of the component, where *k*_*B*_*T* is Boltzmann’s constant times temperature, the thermal energy of the system. In ideal mixtures where the interactions between components can be ignored, the chemical activity is simply proportional to its concentration. This condition is sometimes referred to as the ideal gas limit and is typically valid when solutes are dilute, as is often the case in biochemical measurements of reconstituted proteins. When interactions are relevant, the activity of components can differ dramatically from this linear relationship with concentration. For example, a condition of thermodynamic equilibrium in multi-phase systems is that the μ of every component in each phase is equal. In this case, phases necessarily consist of different concentrations of components but their chemical potentials are equivalent.

In this study, we measure the chemical potential of cholesterol (μ_*c*_) within membranes containing different phospholipid or phospholipid mixtures. Measuring μ_*c*_ in this context provides a new window into the molecular interactions that underlie heterogeneity and phase separation in these well characterized systems. We used methods first described by Ayuyan and Cohen (10) to measure μ_*c*_ in cell membranes by submerging membranes in solutions of methyl β cyclodextrin (MβCD), a sugar that binds cholesterol making it water soluble. These measurements are calibrated against mixtures of ideally mixed cholesterol in an organic phase, making it possible to report μ_*c*_ in units of *k*_*B*_*T*. This enables us to explore quantitative trends, adding to numerous past qualitative measures of cholesterol-lipid interactions in model membranes (11–15). This quantitative measure allows for direct comparison with models of cholesterol-lipid interactions, providing new information towards understanding how these interactions give rise to phase separation and heterogeneity in multicomponent membranes.

## MATERIALS AND METHODS

### Materials

Methyl β-cyclodextrin (MβCD) (CAS: 128446-36-6) was purchased from TCI Chemicals (Portland, OR). Cholesterol (Chol), 1,2-dipalmitoyl-sn-glycero-3-phosphocholine (DPPC), 1,2-dioleoyl-sn-glycero-3-phosphocholine (DOPC), and 1-palmitoyl-2-oleoyl-glycero-3-phosphocholine (POPC) were purchased from Avanti Polar Lipids (Birmingham, AL). N-(7-Nitrobenz-2-Oxa-1,3-Diazol-4-yl)-1,2-Dihexadecanoyl-sn-Glycero-3-Phosphoethanolamine (NBD-PE) was purchased from ThermoFisher (Waltham, MA). Cholesterol Oxidase from Streptomyces was purchased from MP Biomedicals (Santa Ana, CA) and 10-Acetyl-3,7-dihydroxyphenoxazine (Amplex Red) was purchased from Cayman Chemicals (Ann Arbor, MI). All other chemicals and supplies including Horseradish Peroxidase, Raffinose, and Millex-VV Syringe Filter Units, were purchased from Sigma-Aldrich (St. Louis, MO) unless otherwise indicated.

### Preparing MβCD solutions and saturated solutions of MβCD and MβCD/Chol

Solutions containing MβCD and MβCD/Chol were prepared as described previously by other authors (10) with only minor modifications. Briefly, MβCD (5 mg/ml) was dissolved in a buffered saline solution containing 20mM HEPES, 135mM NaCl, 5mM KCl, 1mM MgCl_2_, 1.8mM CaCl_2_, 5.6mM Glucose at pH 7.4. All solutions containing MβCD were degassed for 30 minutes under vacuum and stored under Argon gas to prevent oxidation.

Saturated MβCD/Chol solutions were prepared by first wetting powdered cholesterol (approximately 4mg for a 20mL solution) with 200μL of methanol and drying under Argon followed by 30 min under vacuum to remove residual solvent. This pre-treatment acts to increase the surface area of cholesterol. The 5mg/ml MβCD solution described above was then added to the dried cholesterol, vortexed, then sonicated two times for 2 min each using a Branson bath Ultrasonifier (MODEL S450A, Process Equipment & Supply Inc, North Olmsted OH). The final solution is cloudy with small cholesterol crystals but does not contain large cholesterol aggregates. The saturated cholesterol solution was stored at room temperature with continuous rotation and was typically used at least 12h after preparation. Immediately prior to an experiment, the equilibrated solution is filtered through stacked 0.2 and 0.1μm Millex-VV Syringe Filter Units to remove cholesterol crystals.

The cholesterol standard was prepared by lyophilizing 100μg of cholesterol, then suspending in 10ml of 5mg/ml MβCD buffer. The cholesterol standard was stored under Argon at room temperature with continuous rotation and used for several months.

### Vesicle preparation

Large Unilamellar Vesicles (LUVs) were prepared in glass test tubes pre rinsed with chloroform. Lipid stock solutions in chloroform were combined at the appropriate molar ratios using a Hamilton syringe and included 0.1mol% NBD-PE. Components were then dried under gas while vortexing to form a thin film, then placed under vacuum for 30 minutes to remove residual solvent. The lipid film was hydrated to between 1 and 5mg/ml in an aqueous buffer containing 300mM raffinose and 5mg/ml MβCD, vortexed, then extruded 15 times through a 100 nm Polycarbonate Membrane (part 610005) using mini extruder (part 610000) both from Avanti polar lipids (Birmingham, AL). When DPPC was incorporated, lipids were hydrated and extruded at elevated temperature (>60°C) to prevent phase separation. Typically, vesicle samples used for evaluating μ_c_ contained a final lipid composition of 250μg/ml.

### Measurement of cholesterol and phospholipid concentration

The cholesterol content of aqueous MβCD solutions and suspensions of aqueous MβCD solutions with vesicles were determined using the amplex-red (AR) cholesterol oxidase assay described previously (10) with minor modifications. Stock solutions of cholesterol oxidase (CO; 200U/ml in PBS), horseradish peroxidase (HRP; 200U/ml in Potassium Phosphate buffer, pH 5) were prepared according to manufacturer’s recommendations. AR was stored at 5mg/ml DMSO. The AR reaction buffer was prepared immediately prior to each measurement in the MβCD buffer by adding 1% v/v Triton-X-100, 2U/mL HRP, and 75 μg/ml AR either in the presence or absence of 2U/ml CO.

50μl of each sample was plated in triplicate in a 96 well plate. Plates also included a standard curve prepared from serial dilutions of the saturated MβCD/cholesterol solution in MβCD buffer, a standard curve containing dilutions of LUVs in MβCD buffer, and the cholesterol standard. In some cases, samples were diluted with MβCD buffer prior to mixing with reaction buffer to ensure that readings would fall in a sensitive region of the standard curve. The fluorescence intensity of NBD was then measured using an iD3 Microplate Reader (Molecular Devices, San Jose, CA) with 458nm excitation and emission measured between 510-550nm.

After recording NBD intensities, 50μl of AR reaction buffer was added to wells within the plate. In most cases, plates included 2 technical replicates for each sample with reaction mixture containing CO (CO+) and a single replicate in the reaction mixture lacking CO (CO-). Plates were sealed and incubated for 1h at 37°C, followed by at least 30 min at room temperature prior to recording AR fluorescence intensity with 545nm excitation and emission between 600-650nm.

To determine the effective saturation of cholesterol in samples, fluorescence intensities from CO^−^ wells were subtracted from values obtained from CO^+^ wells on a sample by sample basis. Values corresponding to the standard curve were fit to the following form:

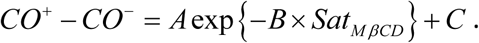

When appropriate, the standard curve was converted to units of cholesterol concentration using a cholesterol standard of 10μg/ml in 5mg/ml MβCD, included in triplicate on every plate. We find that saturated solutions contain 154±25 μM cholesterol when equilibrated at room temperature, which corresponds to a 25±4 MβCD molecules per cholesterol at saturation (Supplemental Figure S1A), in agreement with past measurements (27±3 in 3mg/ml MβCD at 37°C (10)).

To determine the phospholipid concentration of samples, cholesterol concentration was first evaluated in the wells corresponding to the LUV standard curve using the procedure described above. These values were then used to tabulate phospholipid concentration using the known cholesterol mole% of the LUV preparation. We found this step to be necessary because in some cases, it was apparent that LUVs were lost during extrusion and subsequent storage of LUVs. Plots of NBD fluorescence intensity vs. phospholipid concentration were then fit to a line to generate the phospholipid standard curve used to predict phospholipid concentration from NBD intensity in the remaining wells.

In all cases, functions were fit to experimental data points using the fit() function in Matlab (Mathworks, Natick, MA). Errors arising from uncertainty in the standard curve are applied to predicted values using the predint() function within Matlab, using the optional input ‘functional’. When appropriate, errors are propagated to obtain the reported values.

### Calibrating cholesterol activity in MβCD/Chol mixtures

While aqueous MβCD solutions can solubilize some cholesterol, they are complicated solvents and therefore measurements of cholesterol activity in MβCD require calibration. This is accomplished by measuring the saturation of cholesterol in MβCD solutions equilibrated with cholesterol containing hexadecane solutions, of following the protocol described previously (10) with only minor modifications. Cholesterol is expected to mix ideally in hexadecane both because it is a simpler solvent and because the maximum solubility of cholesterol in hexadecane is only a few mol%, meaning that cholesterol concentration remains in the dilute limit even at saturation. As a result, cholesterol activity in hexadecane is expected to be proportional to its concentration. More specifically, the chemical activity of cholesterol within a cholesterol/hexadecane solution is given by its cholesterol concentration divided by the cholesterol concentration of the saturated solution.

Saturated solutions of cholesterol in hexadecane were prepared by dissolving excess cholesterol in hexadecane and equilibrating overnight. The following day, cholesterol crystals were removed by filtration through a 0.45μm syringe filter. This saturated solution, along with a second tube of hexadecane without added cholesterol, were hydrated in excess water (20% of the total volume) and equilibrated for 24h at room temperature under continuous rotation. These solutions were then spun to separate the organic and acquis phases, and the organic phase was retained after centrifugation then filtered through 0.2 & 0.1μm filters. Hydrated hexadecane solutions of a range of cholesterol saturations were assembled by mixing saturated and cholesterol free hydrated solutions at different volume ratios. MβCD solutions over the same range of cholesterol saturations were assembled by mixing saturated and cholesterol free MβCD solutions at the same volume ratios. 500μL of the aqueous MβCD solution was then combined with 100μL of the organic hexadecane solution at each saturation level, capped under argon, and incubated overnight with shaking to equilibrate.

After equilibration, the organic and aqueous phases were separated in each sample through centrifugation. The aqueous phase was retained and transferred into siliconized centrifuge tubes (Sigma-Aldrich Cat. No. T2418) and agitated for 30 min. Solutions were then placed into fresh tubes along with a single polystyrene bead (1/8 inch Polyballs, Cat. No. 17175; Polysciences, Warrington, PA) and agitated again for 30 min to remove trace hexadecane still present in solution. Finally, 50μL of each solution was then transferred to a multiwell plate in triplicate, along with a standard curve containing dilutions of the saturated cholesterol solution in MβCD buffer. Cholesterol content in each well was measured as described above. Aqueous saturation values were normalized to the mean of values from the 100% saturated solution (93%) to account for the small fraction of MβCD whose cholesterol binding is impeded by the small quantities of hexadecane present in acquis solution, following the procedure described in (10). The measured, normalized saturation of aqueous MβCD solutions was then plotted against the constructed saturation of hexadecane solutions they were equilibrated against and fit to the following functional form (Supplemental Figure S1B):

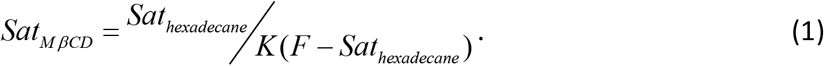

### Evaluating best fit parameters of mean-field models

The following mean-field Gibbs free energies were used to generate expressions for μ_c_ which were then fit to experimental observations. The free energies, implied chemical potentials, and other properties of the models are described at length in the supplementary material. In all cases, μ_c_ was evaluated as an appropriate derivative to match the definition of the chemical potential: 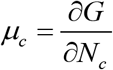. More information can be found in Supplementary Note 1. Several of the following belong to the class of models known as regular solution models. General background for regular solution models is given in Supplementary Notes 2 and 3.

#### Ideal mixing

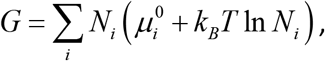

Where N_i_ = N_c_, N_s_, N_u_ is the number of molecules and 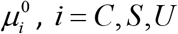 is the chemical potential offset of cholesterol, saturated lipid, or unsaturated lipid respectively. k_B_T is the thermal energy which is given by the Boltzmann constant (k_B_) times temperature (T).

#### Ideal mixing with complexes

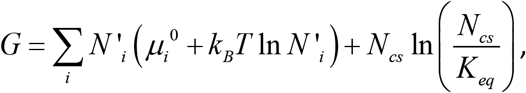

where N’_i_ is the number of molecules of components not found in complexes and the chemical reaction of complex association is: 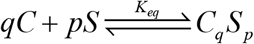. The 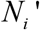 are related to the total numbers of each lipid component by 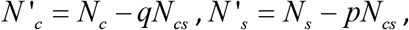, and 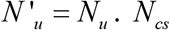 is the number of complexes and is determined by minimizing G as a special case of the procedure described in Supplementary Note 4. We note that a condition for equilibrium is that the chemical potential of cholesterol overall is the same as the chemical potential of cholesterol outside of complexes 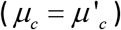.

#### Complexes with interactions

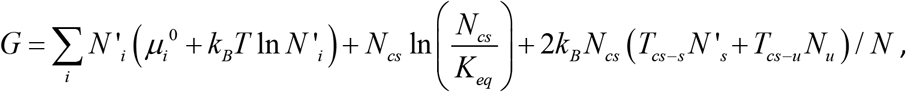

where T_cs-s_ and T_cs-u_ are the critical temperatures associated with complex-saturated lipid and the complex-unsaturated lipid binary systems. All other interaction terms are set to 0. See Supplementary Note 4 for additional information on this model.

#### Regular solution without complexes

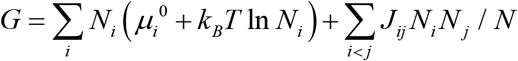

where J_ij_ is the interaction energy between the i^th^ and j^th^ component. Derivations for this model are laid out in Supplementary Note 3.

All fits to μ_c_ were accomplished through a 2 step process to account for the measured uncertainty in both μ_c_ and the mol% of cholesterol. First, a weighted fit was performed where weights were given by the inverse square experimental errors in μ_c_. Errors were then estimated by propagating experimental errors in the cholesterol mol% to μ_c_ using this initial fit, and this contribution to error was added in quadrature to the experimental errors in μ_c_. Finally, a second weighted fit was performed, where weights were given by the updated inverse square error, normalized to average to 1 overall. All fitting was conducted using the fit() function within Matlab. On plots, error bounds are estimated using predint().

### Evaluating approximate phase boundaries and tie lines in mean field models

To determine phase separated regions of the mean field models, we followed the approach of (16). Briefly, the phase separated region is the region of composition space for which the surface *G*(*x*_*C*_, *x*_*S*_) is greater than the convex hull of this surface. For each model, *G* is evaluated using the best fit parameters at a fine grid of compositions (*x*_*C*_, *x*_*S*_), with uniform grid spacing of 2^−10^. The convex hull of the resulting surface is then determined using MATLAB’s convexhull function. The resulting triangulation yields triangles with a long length:width ratio in the phase coexistence region. These long triangles are selected by choosing the triangles where this ratio is more than 20, and the long axis of triangles that meet this criterion are taken as approximate tie lines. The binodal is approximated as the (2-dimensional) convex hull of the endpoints of the tie lines. Finally, the chemical potential within the phase separated region is approximated as a linear interpolation across this region of the chemical potential evaluated at the endpoints of the tie lines.

### Three-component lattice fluid simulations

The regular solution model discussed above corresponds to a mean field approximation to a three-component lattice fluid with nearest-neighbor interaction Hamiltonian given by

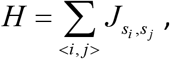

where < *i, j* > indicates that the sum is to be taken over nearest neighbors, *s*_*I*_ is the component at site i, and *J*_*kl*_ is the interaction energy between species k and species l. To compare to the full thermodynamics of this system, it is simulated using home-built C and MATLAB code implementing a fixed-composition Metropolis sampler for this Hamiltonian.

The fixed composition criterion is enforced by only proposing updates that swap the components at two sites. Unless otherwise specified, the simulations are performed on a periodic LxL site square lattice with L=256, with 1000xLxL swaps proposed between samples, and the first 50 samples are discarded as burn in.

For each simulated condition, μ_*C*_ is obtained from 7 simulation samples following a procedure discussed in detail in Supplementary Note 5. In brief, we define

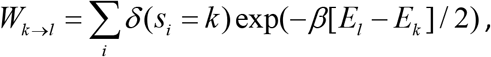

where *k, l* index chemical species, *i* indexes sites of the lattice, *s*_*i*_ is the species at site *i*, and *E*_*k*_ is the value of the Hamiltonian with site *i* replaced by species *k*. By reference to a Monte Carlo scheme exchanging lipids between the simulated lattice and a bath of particles at specified chemical potentials μ_*j*_, *j* = *C, S,U*, it may be shown that

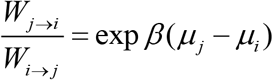

when the lattice is in equilibrium with the bath. The calculated chemical potential differences are integrated over composition space to estimate the overall free energy per molecule g = G/N. Finally, μ_*C*_ can be obtained from *g* and the chemical potential differences using the identities derived in Supplementary Note 1.

To determine the phase diagram implied by the simulations, tie lines are estimated from individual samples of each simulated composition. Local composition images are computed by convolving the binary matrix of site occupancies for each component with a disc with a diameter of 25 lattice sites. For a phase-separated composition, 2d histograms of these local compositions with respect to Cholesterol and DPPC mole fractions show two peaks, with compositions equal to those of the ends of the tie-line that the overall simulated composition falls on. Compositions that do not phase separate are characterized by a single peak in their local composition histograms.

## RESULTS

### Measurements of cholesterol activity in cholesterol phospholipid mixtures

Cholesterol activity in vesicles was determined by measuring the cholesterol saturation in MβCD solutions in contact with LUVs with different cholesterol and phospholipid contents at room temperature as illustrated in Figure 1. This was accomplished by mixing LUVs of defined phospholipid and cholesterol composition in MβCD/cholesterol mixtures with a range of initial cholesterol chemical activities. After an equilibration period to allow for cholesterol to exchange between vesicles and MβCD in the aqueous phase, some of the equilibrated sample was retained for later analysis and the remaining suspension was pelleted through centrifugation (18k × g for 90 min @23°C). The majority of the supernatant was then extracted to produce a solution that was depleted in vesicles, while the pellet and remaining liquid were mixed to produce a solution enriched in vesicles (Figure 1A). To facilitate a robust depletion of LUVs from the supernatant, LUVs were prepared by hydrating lipids in a solution of 5mg/ml MβCD and 300mM raffinose, osmotically matched to the MβCD containing solutions used to manipulate and measure cholesterol activity. Cholesterol and phospholipid concentrations were then determined in the initial vesicle suspension, the pellet and the supernatant samples. We note that past studies have separated MβCD solutions from LUVs through filtration (10, 11, 13). In our hands, we found that not all cholesterol could be accounted for using this protocol when vesicles were present, suggesting that some remained adhered to the filter. We also note that although past work has measured phospholipid extraction from vesicles by high concentrations of MβCD, the concentrations we use for these experiments lead to negligible phospholipid extraction (13). Therefore, phospholipid extraction by MβCD is not expected to be a substantial source of error in our measurements.

**Figure 1:**
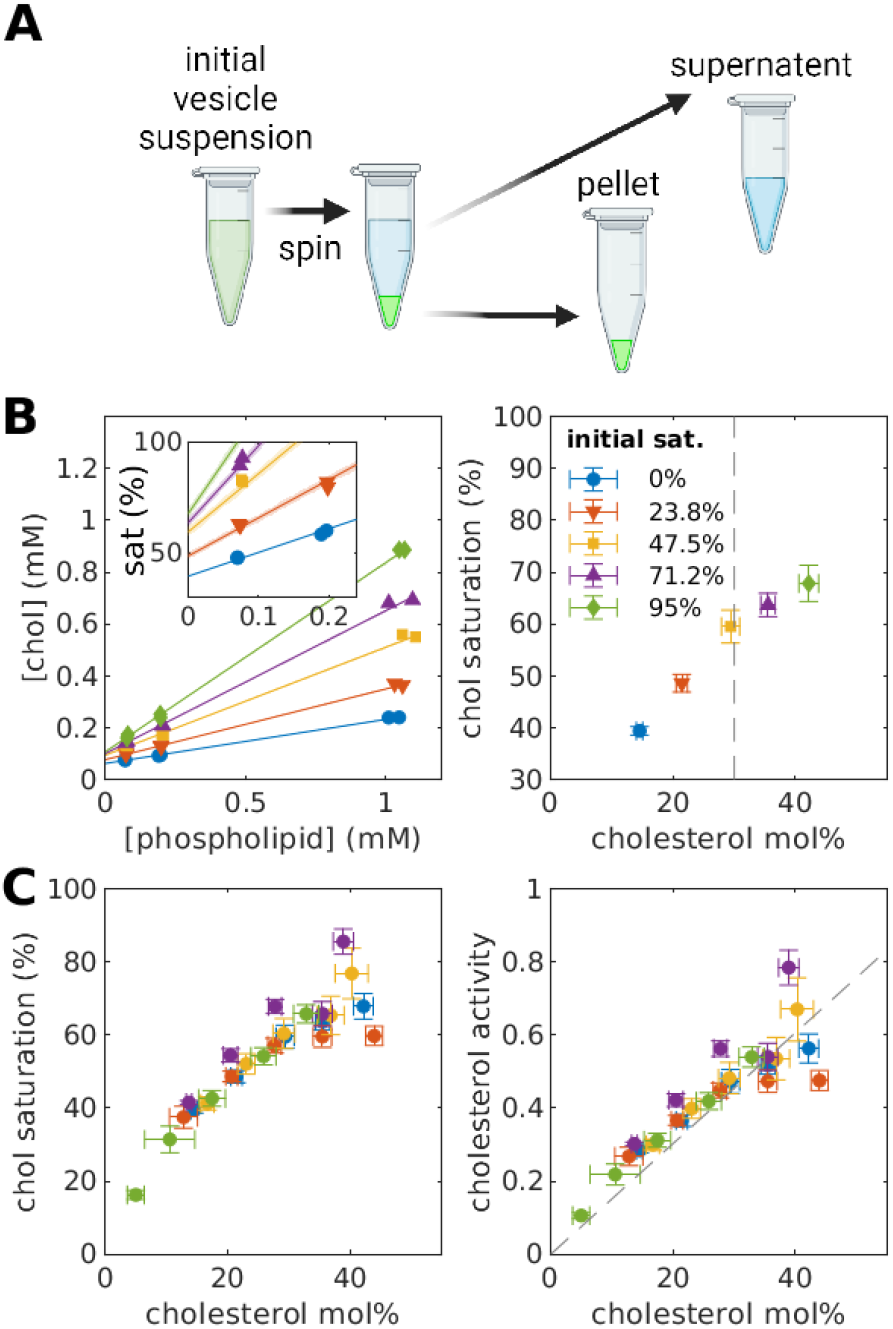
Experimental determination cholesterol activity in vesicle membranes. A. Schematic representation of the measurement scheme. Vesicles were suspended in MβCD/cholesterol solutions of varying initial cholesterol activities. After an incubation period, suspensions were spun to generate samples with varying enrichment of phospholipids. B. Cholesterol and phospholipid concentrations were measured for supernatant, pellet, and initial suspensions including technical replicates (left). This example used vesicles of 70% DOPC and 30% cholesterol initially suspended in MβCD/Chol solutions with the saturations shown. A linear trend is fit to extract the y intercept (cholesterol saturation in the absence of vesicles), and the slope (cholesterol to phospholipid ratio in vesicles), which are plotted for each sample (right). C. The results of multiple trials with different DOPC/Chol vesicle preparations collapse into a single curve that can be plotted vs. cholesterol saturation (left) or converted into cholesterol activity (right) using the calibration curve of Sup Fig S1B.

While the centrifugation protocol described above did not fully separate LUVs from MβCD/cholesterol solutions, both the concentration of cholesterol in solution and the mole fraction in vesicles could be deduced. Figure 1B shows a representative measurement for DOPC vesicles initially containing 30 mol% cholesterol to illustrate the approach. First, values corresponding to phospholipid concentration and cholesterol saturation are plotted for each measured solution. In this example, different colored points originate from suspensions containing different initial cholesterol saturation levels. Points clustered around different low, intermediate, and high phospholipid concentrations originate from the supernatant, initial, and pellet samples respectively. Points corresponding to the same initial suspensions are then fit to a line. The y-intercept of this line represents the cholesterol saturation of the MβCD solutions in the absence of phospholipid (inset), while the slope indicates the cholesterol to phospholipid ratio of vesicles, which is algebraically related to cholesterol mol%. These values are plotted with associated error bounds for this single experiment in Figure 1B. Multiple measurements were carried out on different days, sometimes with different initial vesicle cholesterol concentrations, and results fall along the same curve (Figure 1C). Cholesterol saturation in MβCD is then converted to cholesterol activity using the calibration curve from Supplementary Figure 1.

### Measurements of μ_c_ in binary mixtures with DOPC, POPC, and DPPC

Figure 2A shows how cholesterol chemical activity varies with cholesterol fraction in vesicles of different phospholipids, highlighting the significant impact of phospholipid chains. Previous studies have found a maximum cholesterol solubility of 67% in several membrane systems (17–19). With fully saturated solutions defining activity 1, this implies that ideal mixing in these systems would correspond to a straight line when plotting chemical activity vs concentration, connecting the origin to the point with 67% concentration, activity 1 (dashed line in 2A). Our data in DOPC membranes are close to this prediction without introducing fit parameters, implying weak interactions between cholesterol and DOPC, a lipid that contains two mono-unsaturated oleic acid acyl chains. In contrast, cholesterol activity remains low in mixtures with DPPC until close to 40 mol% cholesterol, a strong deviation from ideal mixing. DPPC contains two saturated palmitic acid acyl chains, and our observations reflect the presence of stronger attractive interactions between cholesterol and DPPC. POPC contains both palmitic and oleic acid chains, and cholesterol activity takes on intermediate values when incorporated into POPC LUVs. We note that DPPC/Cholesterol mixtures are known to separate into liquid-ordered and solid phases below 30% cholesterol (8), likely contributing to why the activity remains low even at elevated cholesterol concentrations. We found that DPPC membranes prepared with 20% or 30% cholesterol did not equilibrate with MβCD-cholesterol mixtures at room temperature, possibly due to the presence of gel or solid phase in these membranes. All points shown for DPPC initially contained 40% cholesterol and equilibrated for at least 24h. Also shown on Figure 2A are measurements replotted from past measurements of red blood cell (RBC) membranes at 37°C (10). Cholesterol activity in these cell membranes are far from the expectations of ideal mixing and more closely trend with measurements of POPC and DPPC in different cholesterol concentration regimes.

**Figure 2.**
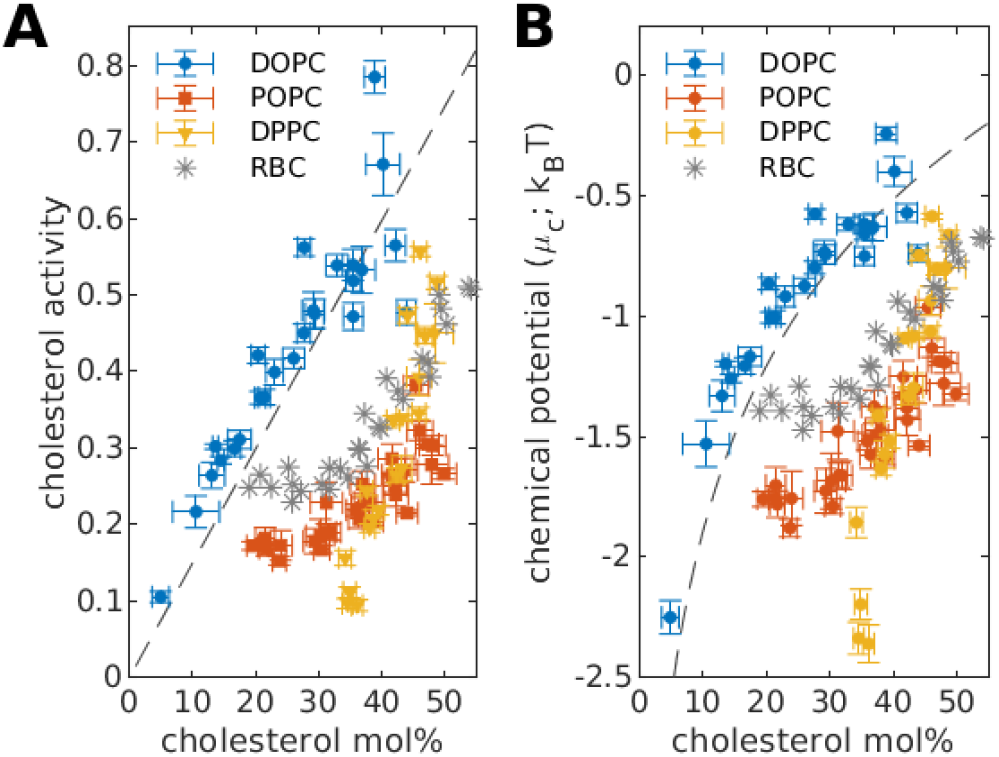
Cholesterol activity (A) and chemical potential (B) in binary PC/Chol membranes. The dashed lines indicate expectations of ideal mixing, assuming a maximum solubility limit of 67% cholesterol in PC membranes. The gray points are measurements in red blood cells (RBC) replotted from (10).

When cholesterol chemical activities are converted to chemical potentials (μ_c_), we find that μ_c_ takes on values between −2.5 and −0.5 k_B_T for the cholesterol/phospholipid mixtures investigated (Figure 2B). The relative trends apparent in Figure 2 are in good agreement with a large number of past studies where properties related to chemical potential have been measured but not calibrated to obtain values in units of energy (11–15).

### Measurements of μ_c_ in ternary mixtures of DOPC, DPPC, and cholesterol

We have also measured μ_c_ in vesicles containing both DOPC and DPPC, and results are summarized in Figure 3. In these measurements, phospholipids were prepared at a specific molar ratio with cholesterol, then vesicles were incubated in MβCD/cholesterol solutions as in Figures 1 and 2. It is assumed that vesicles remain intact throughout the MβCD/cholesterol incubation, resulting in vesicles that retain a fixed molar ratio of DOPC to DPPC but with varying cholesterol content. As before, the results of measurements from different trials collapse onto a single curve for each DOPC/DPPC ratio interrogated (Figure 3A), although in some cases there is significant scatter of points. To smooth noise, results for specific DOPC/DPPC ratios were fit to polynomials to capture both the trend and the confidence interval of the measurement (solid lines and shaded regions in Figure 3A respectively). These fits are used to interpolate the chemical potential surface shown in Figure 3B.

**Figure 3:**
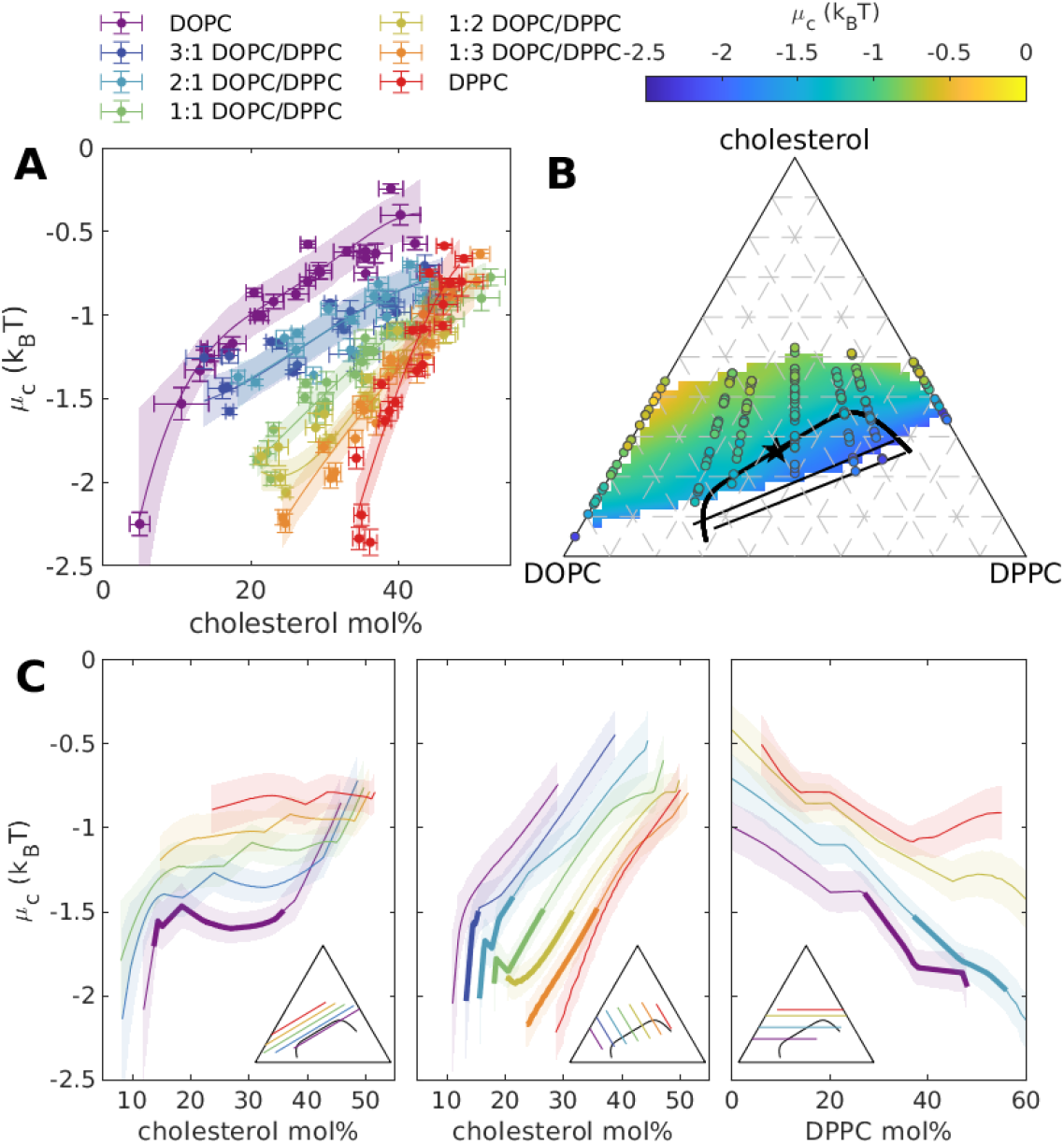
μ_c_ in ternary DOPC/DPPC/cholesterol membranes. (A) Measured μ_c_ for vesicles prepared with the specified DOPC/DPPC ratios. Measurements were taken over several days with multiple vesicle preparations. Solid lines are fits to polynomial functions and shaded regions indicate 68% confidence intervals of predicted values. (B) The μ_c_ surface observed at room temperature extrapolated from the fits shown in part A with measured points indicated as circle symbols. Black straight lines are tie-lines redrawn from (20) for DOPC/DPPCd62/Chol at 17.5°C to account for differences in the melting temperature between protonated DPPC and deuterated DPPCd62. The curved line is an estimate of the miscibility phase transition boundary (binodal), rotated slightly from the one reported at 17.5°C in (20). (C) Trajectories through the extrapolated μ_c_ surface in the direction of tie-lines (left), perpendicular to tie-lines (middle) and at constant cholesterol (right). Shaded regions indicate 68% confidence intervals and compositions of the individual cuts are indicate on the triangles shown as insets.

Mixtures of DOPC/DPPC/Chol undergo liquid-liquid and solid-liquid phase separation, and Figure 3B also includes tie-lines measured for a closely related lipid mixture by deuterium NMR (20) and an estimated miscibility gap. We note that the phase diagram here is expected to be slightly different than the one presented in this past work, both because the protonated DPPC lipid is used, and because a small mole% of NBD-PE is included (0.1%) (21). The measured μ_c_ surface includes regions both inside and outside of the miscibility gap but does not extend to compositions where gel or solid phases are reported at lower cholesterol and higher DPPC concentrations.

Figure 3C shows linear trajectories through the measured μ_c_ surface. These include trajectories that run in the direction of tie-lines within the miscibility gap (left). As expected, μ_c_ remains constant along the tie-line within the coexistence region, a requirement for chemical equilibrium in phase-separated systems (22). Trajectories that run parallel to tie-lines but pass outside of the miscibility gap also retain a shallow slope over the range of compositions interrogated. Trajectories through the μ_c_ surface that run perpendicular to tie-lines increase with increasing cholesterol concentration (center). No obvious discontinuities in slope are observed as the phase boundary is crossed, but this may be due to the smoothing process employed or the sparse sampling of data points in some regions. Finally, trajectories at constant cholesterol concentration are shown (right), highlighting the non-ideality of these ternary mixtures. Ideal mixtures have chemical potentials that depend only on the concentration of the component. In contrast, these constant cholesterol trajectories show that μ_c_ varies by several k_B_T as phospholipid ratios are varied.

### Measurements of μ_c_ limit the binding affinity of cholesterol-phospholipid complexes in a condensed complex model

μ_c_ is the derivative of the Gibbs free energy with respect to the number of cholesterol molecules and is therefore specified by thermodynamic models. Figure 4A fits experimental observations of μ_c_ to a thermodynamic model with no interactions, in which the chemical activity of cholesterol is simply proportional to its concentration. This model has a single fit parameter, 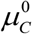, which defines the solubility limit and therefore proportionality constant. As expected, this simple model does not capture important features of the experimental data, most notably the dependence of μ_c_ on the DOPC/DPPC ratio. In this formulation, cholesterol has no information about non-cholesterol species; there is only cholesterol and not-cholesterol. In addition, the best fit parameter value for μ_o_ is unphysical because it corresponds to a maximum solubility limit of 110% cholesterol.

**Figure 4:**
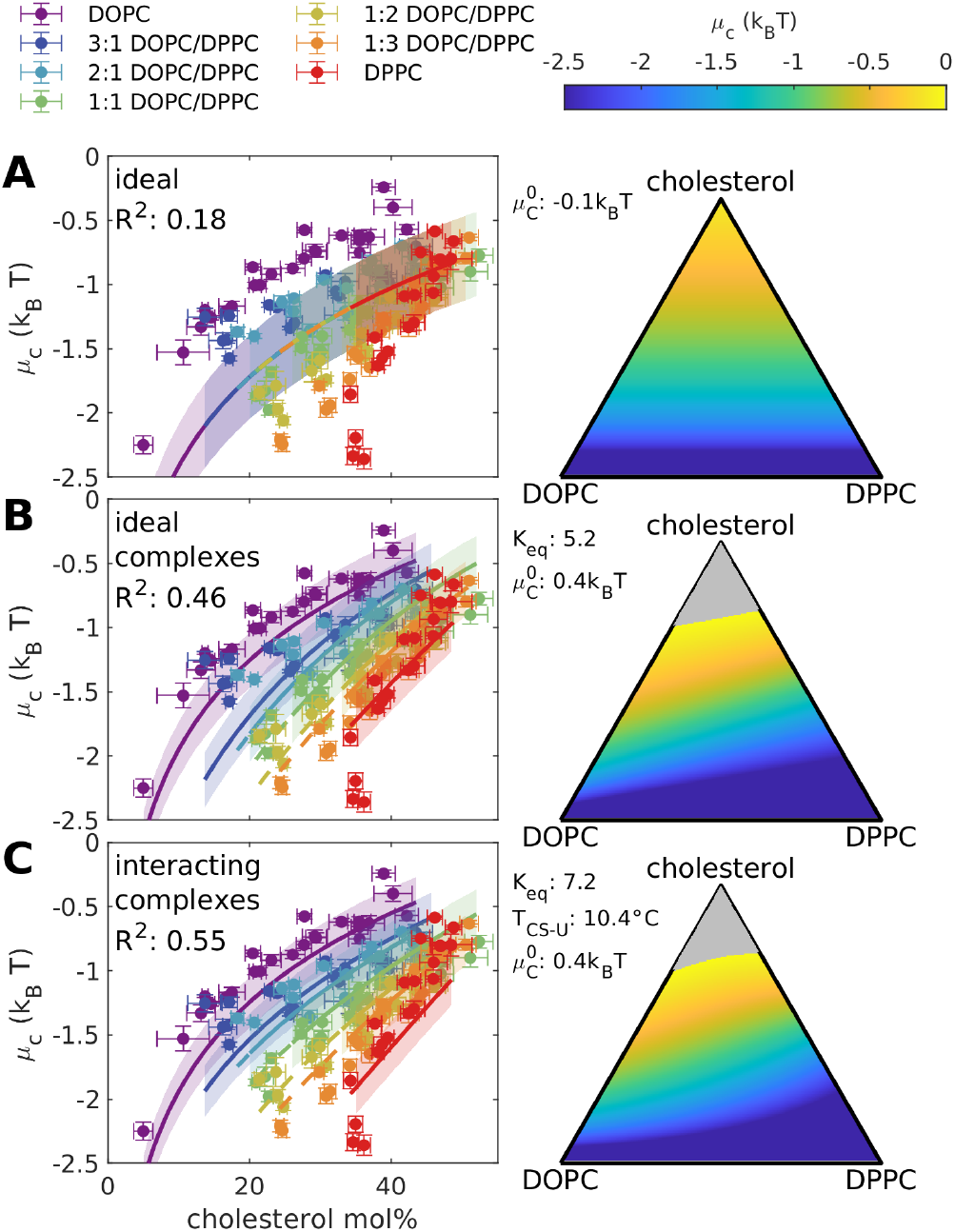
Fits of μ_c_ to ideal mixing (A) and two models with condensed complexes (B, C). (left) Experimental measurements of μ_c_ replotted from Figure 4 (points) are simultaneously fit to the indicated model as described in Methods. The best fit solutions are plotted along each DOPC/DPPC series (lines) with predicted error bounds indicated as shaded regions. (right) Surface plots of the best fit solution extrapolated to all values in the Gibbs triangle. Gray regions indicate compositions beyond the solubility limit, i.e. μ_c_>0. Best fit parameter values are indicated, as is the Pearson’s R^2 value at the best fit. Confidence intervals on best fit parameter values are provided in Sup. Table 1.

Previous work has proposed a condensed complex model to describe aspects of phospholipid-cholesterol phase diagrams in monolayer and bilayer membranes (23–25). In this model, cholesterol and phospholipids assemble into complexes of fixed stoichiometry with an affinity characterized by its dissociation constant K_eq_. Condensed complexes can additionally interact with saturated and unsaturated lipids to produce phase separation. Figure 4B presents a model that retains ideal mixing but allows for the formation of a condensed complex made up of cholesterol and DPPC with a fixed stoichiometry of 1:1, as done in past work (24, 25). The model used in Figure 4B contains 2 parameters, 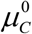 and the equilibrium constant of complex formation (K_eq_). This model produces much better agreement with experimental measurements of μ_c_ than the ideal mixing model of Figure 4A. Shifts towards larger cholesterol mol% for smaller DOPC/DPPC ratios are captured by this model, and the best fit μ_o_ corresponds to a solubility limit of 69% chol in DOPC and 75% in DPPC. The best fit value for K_eq_ is 5.2, corresponding to a binding affinity of 1.6 k_B_T. This low association energy means that the fraction of cholesterol not bound in complexes remains substantial across all lipid compositions.

The models of Figure 4A,B do not support phase separation by construction, since the Gibbs free energy does not include interactions between components. Figure 4C fits the measured μ_c_ to a model that includes both complex formation and repulsive interactions between complexes and DOPC. The model used in Figure 4C contains 3 parameters, 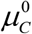 and K_eq_ as in Fig 4B, and an interaction energy, represented as a critical temperature for phase separation between complexes and unsaturated lipid (T_CS-U_). Including this interaction allows for a better fit to experimental measurements, but the fitted parameters are not consistent with a phase separating mixture at room temperature. Here again, the fitted equilibrium constant K_eq_ is small (K_eq_ = 7.2), indicating weak binding of this complex. We also considered other elaborations of this class of model, including one where the interaction energy between the complex and DPPC was allowed to vary (*T*_*CS* −*S*_ ≠ 0), and a second with different complex stocheometery, as shown in Supplementary Figure S2-3. In both cases resuts were either not improved compared to the models of Figure 4 or were deemed unphysical.

### Measurements of μ_c_ are described by models capturing pairwise interactions between components

We additionally investigated whether experimental observations could be described by models that do not allow for the formation of complexes, but instead include pairwise interactions between all three components (26). Two approaches are shown in Figure 5. In the first, we employ a mean field regular solution theory model of the ternary system and the best fit solution is shown in Figure 5A. There are 4 parameters in this model: μ_o_ and interaction energies between all three pairs of components, between saturated and unsaturated (J_S-U_), saturated and cholesterol (J_C-S_) and unsaturated and cholesterol (J_C-U_). This model accurately captures trends in measured μ_ch_. This model also supports phase separation, and the binodal line and several tie lines are shown. Here, phase separation is predicted over a range of compositions in qualitative agreement with past experimental observations. This model does not incorporate interactions that would give rise to a solid phase, therefore the phase diagram does not capture regions of solid-liquid coexistence or the 3 phase region detected experimentally (20).

**Figure 5.**
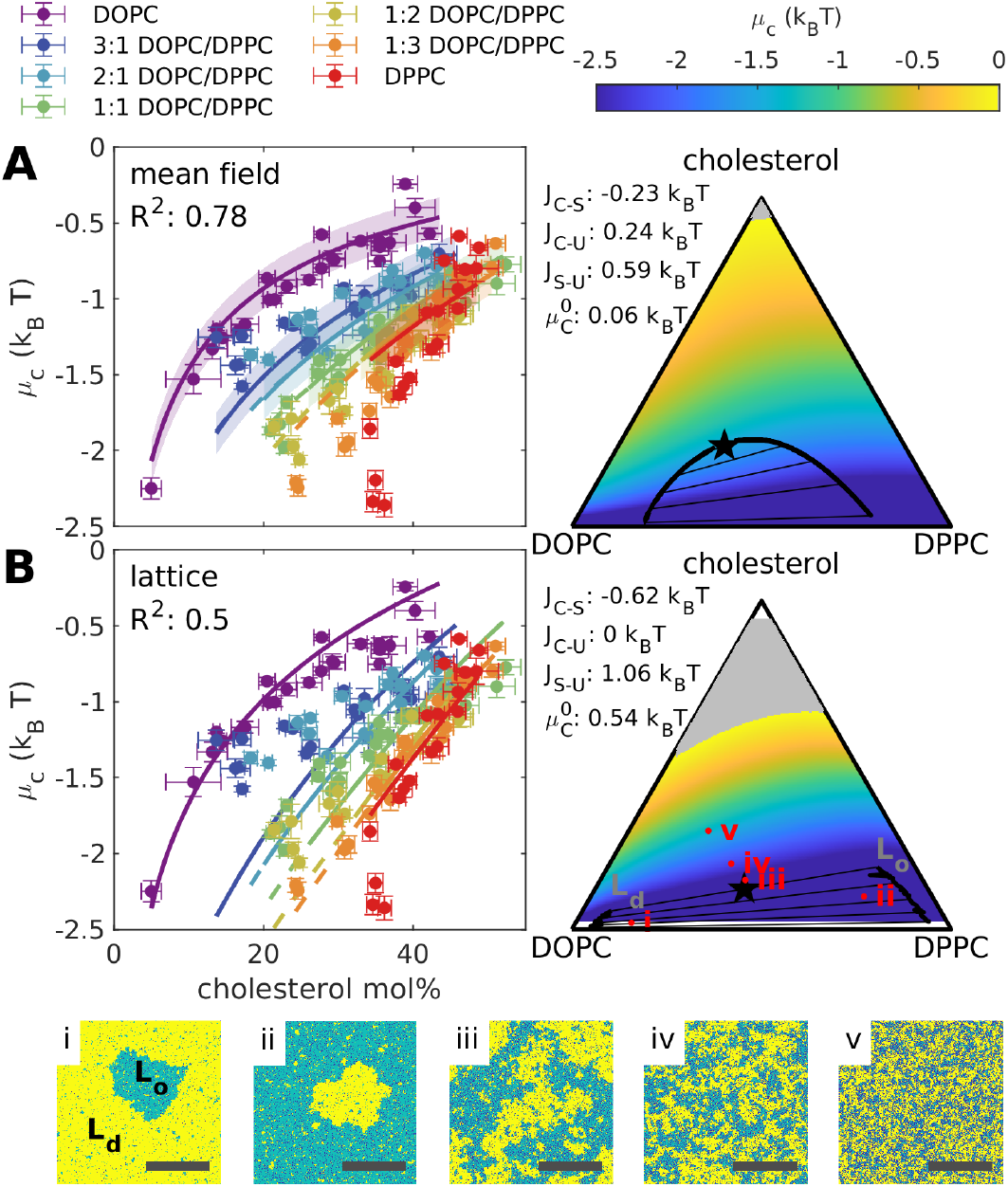
Fits of μ_c_ to mean field (A) and lattice (B) models of pairwise interactions between components. (A) (left) Experimental measurements of μ_c_ replotted from Figure 4 (points) are simultaneously fit to the regular solution theory mean field model as described in Methods. The best fit solutions is plotted along each DOPC/DPPC series (lines) with predicted error bounds indicated as shaded regions. Dashed lines indicate conditions within the 2-phase region. (right) Surface plot of the best fit solution extrapolated to all values in the Gibbs triangle. (B) (left) Experimental measurements of μ_c_ (points) and interpolated values extracted from lattice model simulations (lines). (right) Surface plot of interpolated μ_c_ values evaluated over multiple simulations. Several simulation snapshots are shown (Bi-v), corresponding to the compositions marked on the Gibbs triangle. Representative “Lo-like” and “Ld-like” compositions are indicated in (i). Scale bar: 100 lattice spacings. (A, B) Gray regions indicate compositions beyond the solubility limit, i.e. μ_c_>0. Regions of phase separation are evaluated from snapshots as indicated by the binodal (thick black line) and tie-lines (thinner lines) that merge at a critical point (star). Best fit parameter values are indicated, as is the Pearson’s R^2 value at the best fit. Confidence intervals for fitted parameters are provided in Sup. Table 1.

The analytical models described in Figure 4 and 5A are mean-field models that consider average effective concentrations. Membranes, which exist in two dimensions (2D), are expected to exhibit deviations from this mean-field approach, since the mean field approach ignores fluctuations whose probability is not negligible. These differences are the most striking close to the critical point of these systems, where in particular the shape of the phase boundary differs between 2D and mean-field predictions (22). To address this, we additionally conducted simulations of pair-wise interactions between components on a 2D square lattice and extracted μ_c_. Unlike the analytical models of previous figures, estimation of the Gibbs free energy G must be carried out by numerical integration of its derivatives with respect to composition, so accurate estimates depend on performing Monte Carlo simulations at many compositions. Our estimates of μ_c_ depend on this computationally costly estimate of G, so are not amenable to nonlinear fitting to determine the most probable parameter coefficients. Instead, we first sought to find values of J_C-S_ and J_C-U_ that produced values of μ_c_ that were consistent with experimental trends in the limiting binary mixtures at the same μ_o_. Then, the remaining parameter J_C-S_ was estimated by conducting a series of simulations at fixed cholesterol fraction considering a range of J_S-C_ values. Finally, simulations of the full model over the entire composition space were conducted for a range of parameter values close to these initial estimates. The model shown in Figure 5B represents the model that yielded the best agreement with experimental observations. Of note, this model contains a coexistence region that terminates in a critical point at 12 mol% chol. Several representative simulation snapshots also are shown.

Contrasting the mean-field and lattice-model approaches of Figure 5, we find that the mean-field approach is more successful at capturing trends in μ_c_ for mixtures with high DOPC/DPPC ratios (blue points) whereas the lattice model is more successful at capturing trends in in μ_c_ for mixtures with low DOPC/DPPC (red points). This is likely related to the shape and extent of the miscibility gaps, which are largely dictated by the models themselves. In both cases, the model dictates that binodals are symmetric around 1:1 DOPC/DPPC in the absence of cholesterol. In real membranes, the presence of additional interactions likely breaks this symmetry, allowing for a broader range of possible compositions for the coexisting phases. As cholesterol is increased, tie-lines tilt due to the presence of more favorable interactions between cholesterol and DPPC vs. DOPC. Both models contain a critical point although the shape of the binodal near this point differs dramatically between the two models, as is expected due to their different critical exponents.

## DISCUSSION

Cholesterol is vital for a broad range of cellular functions, and extensive past work demonstrates its multifaceted roles in tuning the physical properties of membranes. In this study, we have measured the chemical potential of cholesterol in purified binary and ternary membranes, following protocols developed for measuring μ_c_ in cells. μ_c_ is a derivative of the Gibbs free energy, therefore its measurement provides a window into the thermodynamics of these systems. Moreover, the activity of a component, rather than its concentration, reflects its availability, for example to bind receptors.

### Using MβCD as a tool to study membrane thermodynamics

Thermodynamic studies of bilayer membranes are typically conducted under conditions of fixed composition, with individual vesicles having a predetermined ratio of components. This is because the solubility of lipids in aqueous buffers is so low that there is minimal exchange between membranes on experimental time-scales. Incorporating MβCD in solution changes this situation, enabling cholesterol molecules to rapidly exchange between vesicles and the aqueous phase. Under this condition, the molar fraction of cholesterol will adjust in vesicles to minimize the free energy of the entire vesicle suspension. In one regard, this makes the experiment more difficult because we are required to measure the mol% of cholesterol in vesicles at the end of the experiment. Alternately, this experimental modality gives us access to new information, namely the chemical activity of cholesterol at equilibrium. When combined with knowledge of the phase diagram from experiments at fixed composition, this can put limits on the free energy of the system.

At equilibrium, the chemical activity of the same component in different states must be equal. This condition for equilibrium is exploited to infer the cholesterol activity in MβCD-containing aqueous solutions as a function of their saturation by equilibrating solutions with cholesterol dissolved in hexadecane. This condition of equilibrium is also exploited to infer the cholesterol activity in membranes by measuring cholesterol activity of MβCD solutions in contact with membranes. The combination of these measurements allows for the quantitative measurement of μ_c_. The application of this elegant approach (10) depends only on our ability to maintain chemical equilibrium, a constraint that factored into our experimental strategy.

MβCD solubilizes cholesterol but is very inefficient. We find that it takes 25±4 MβCD molecules to solubilize a single cholesterol molecule at saturation, using commercially available MβCD stocks (Supplementary Figure S1). This suggests either that collective behaviors of MβCD molecules are required to solubilize cholesterol, or that only a small subset of MβCD molecules are capable of cholesterol binding. In support of the first possibility, cyclodextrins including MβCD are known to assemble into higher order structures (27, 28). This could explain why MβCD/cholesterol mixtures do not robustly cross spin filters or dialysis membranes even when high molecular weight cutoffs are used. Separate studies document the importance of methylation in controlling MβCD/cholesterol binding (29). The commercial MβCD reagent used in the current study claims to contain a broad distribution of methylation levels, leaving open the possibility that only a small subset of MβCD monomers can bind cholesterol. Whatever the cause, improved solubility of cholesterol in MβCD would reduce noise and improve the dynamic range of our measurements by equalizing cholesterol concentration between the vesicle and soluble pools.

Despite experimental limitations, we measure robust trends in μ_c_ across experiments, providing support for the validity of our approach. Notably, μ_c_ generated by combining data from a wide range of individual measurements independently verifies known aspects of the phase diagram of this system, namely tie-lines map out regions of constant μ_c_ (Figure 3C). Moreover, the quantitative trends reported here are in good agreement with qualitative observations in previous studies that measured the equilibrium partitioning of cholesterol between vesicles and MβCD solutions, or between vesicles of different lipid compositions facilitated by MβCD (11–14). These past measurements demonstrate that cholesterol partitions more strongly into membranes with greater lipid saturation.

### Measurements of μ_c_ reinforce that cholesterol containing membranes are not ideally mixed

In well mixed systems, the chemical activity of components is simply proportional to the concentration of that component. The measurements presented in Figs 2 and 3 provide many instances that violate this simple picture. For one, μ_c_ is dependent on the identity of other lipids in the membrane, meaning the same cholesterol mol% produces vastly different cholesterol activities depending on which other components are present. For the ternary DOPC/DPPC/cholesterol membrane, this non-ideality is apparent when taking constant cholesterol slices through the μ_c_ surface, where it is observed that μ_c_ can vary by several k_B_T. This is also apparent when examining that μ_c_ remains constant over a range of cholesterol concentrations when traversing a tie-line in the miscibility gap. While this is expected, it is striking that this behavior persists well beyond the limits of the phase separated region, emphasizing that the same interactions that give rise to phase separation also impact the physical properties of membranes outside of phase separation.

This non-ideal mixing means that the activity of cholesterol depends not only on cholesterol’s concentration, but also on how favorable its interactions are with other membrane components. These effects can be quite large. For example, in binary mixtures of DOPC and 30 mol% cholesterol, μ_c_ is almost 2K_B_T larger than in binary mixtures of DPPC and 30 mol% cholesterol, corresponding to a more than five-fold difference in chemical activity. This particular example highlights pitfalls associated with equating concentration and chemical activity in multicomponent membranes because an identical concentration of cholesterol is much more (>5x) available to bind proteins when incorporated into a DOPC membrane than when incorporated into a DPPC membrane. While results in these model systems do not extend directly to biological membranes, they do highlight the possibility that processes which require cholesterol can be regulated by modulating the chemical activity of cholesterol without changing its concentration.

### Measurements of μ_c_ provide complementary information to the phase diagram

Both the chemical potential and the phase diagram can be inferred from the Gibbs free energy. The chemical potential of a component is given by the derivative of the Gibbs free energy with respect to that component’s particle number. This means that, if we could measure the chemical potential of all three components, we could in principle integrate these to infer the Gibbs free energy in the space of possible compositions, up to a constant of integration. This would be sufficient to infer the phase diagram and all thermodynamic properties.

With a measurement of just a single component’s chemical potential we cannot unambiguously infer the phase diagram. However, we can still see signatures of phase coexistence in these measurements. For example, in paths taken along tie-lines μ_c_ remains constant even as the concentration of cholesterol changes. Moreover, deviations of the chemical potential from ideal gas predictions reflect the influences of the same interactions that do sometimes lead to phase coexistence, even in regions of composition space, or temperatures without phase separation.

### Experimental trends in μ_c_ can be attributed to weakly bound condensed complexes

To understand our measurements in DOPC/DPPC/Chol membranes, we used multiple theoretical and simulation approaches. We note that none of these models are constructed to replicate interactions that give rise to regions of solid-liquid coexistence, as is experimentally observed in this system at high DPPC and low cholesterol compositions, therefore we anticipate disagreement between models and experiment in this regime.

We first modified a model introduced by Radhakrishnan and McConnell (23–25), in which cholesterol forms condensed complexes with saturated lipids with fixed stoichiometry. In the first iteration, we allow for the formation of complexes but do not include additional interactions between lipids or lipids and complexes. In the absence of additional interactions, the chemical activity can be simply interpreted as the concentration of cholesterol that is not bound within complexes. This model is moderately successful in capturing experimental trends in μ_c_ but does not phase separate by construction. Including interactions between complexes and DOPC (25) allows for better agreement with measurements of μ_c_, but we find that the fitted interaction strengths are not consistent with room-temperature phase separation of these mixtures as is observed experimentally (20, 30).

Past applications of this model assumed that complexes were tightly bound with a large equilibrium constant of K_eq_ close to 150 (24, 25). Here we find that measured chemical potentials are only consistent with this condensed complex model if the complexes are relatively weak, with a dissociation constant >20 times lower than previously assumed (K_eq_ ~ 5-7 in Figure 4B-C). Our measurements suggest that complexes, if present, are held together by interactions that are on the order of the thermal energy (<2k_B_T). One consequence of this observation is that individual cholesterol molecules would be expected to readily exchange between complexed and free forms on experimental timescales, providing opportunities for enzymes such as cholesterol oxidase to act on unbound cholesterol even if a significant fraction is bound.

### Experimental trends in μ_c_ can be described by pair-wise interactions between components

We also have presented both mean field and lattice models where deviations from ideal mixing come from interactions between cholesterol molecules and their neighbors. Here again we find that models capture experimental trends in μ_c_. In addition, the model parameters that best explain the μ_c_ measurements also reproduce the broad qualitative shape of the experimental phase coexistence region for this lipid mixture. Some features of the coexistence region are poorly matched to experiment, such as the width of the coexistence region. We note that other groups have observed similar difficulties when fitting three-component models of this type to phase diagrams of lipid mixtures (16, 31). In addition, others have reported extensions to this class of model that have proven successful in fitting other types of experimental data from few-component lipid mixtures, including mixtures with cholesterol (1, 16, 32). It is clear that a full explanation of all available experimental data for these mixtures would require a substantially more detailed model.

While these models lack explicit complexes, fitted parameters imply that cholesterol has favorable interactions with DPPC, meaning that it will have a lower activity when cholesterol molecules tend to be surrounded by DPPC lipids. In this sense, while complex formation brings a different microscopic picture to mind, it leads to similar thermodynamic predictions in the limit where complexes are relatively weak.

### Comparison to past results in model membranes and cells

μ_c_ is a physical observable reflecting both the concentration of cholesterol and how energetically unfavorable it is to remove cholesterol molecules from their local lipid environment. While past studies have been concerned with comparing and matching the concentration of cholesterol to physiological values in model membranes, our results stress that it may also be important to match cholesterol’s activity. Directly measuring μ_c_ in calibrated k_B_T units provides a robust means to compare measurements across experiments and systems. A comparison of past results in red blood cell membranes (10) and current results in model membranes is shown in Figure 2, where it was found that RBC membranes more closely resemble POPC and DPPC membranes than they do DOPC membranes. The non-ideality of RBC membranes is striking, with low cholesterol activity even when membranes contain a large mol% of cholesterol. This indicates that cholesterol experiences attractive interactions with membrane components, which in the case of RBC membranes could involve both proteins and lipids. Past work also measures relatively low cholesterol activity in two mammalian cell culture lines (between −1.4 to −2.2 k_B_T) (10). These values correspond to <15 mol% cholesterol in DOPC membranes or less than 40 mol% cholesterol in DPPC membranes, which is striking because cells in culture typically contain 30-40 mol% cholesterol and a large fraction of unsaturated and polyunsaturated lipids (33). One important difference between the model membranes studied here and cell membranes measured previously is the presence of phospholipid and cholesterol asymmetry across leaflets. Recent reports suggest that cholesterol plays a role in supporting this asymmetry (34, 35), and it is possible that the surprisingly low μ_c_ detected for cells is related to this phenomenon.

There is a body of past work that reports cholesterol chemical activity inferred in model membranes and cells from kinetic measurements of cholesterol extraction or oxidation with MβCD or cholesterol oxidase respectively (11, 17, 36, 37). These past studies argue that initial rates are proportional to the chemical activity in the equilibrium state. While this measurement almost certainly correlates with the activity of cholesterol, it could also be impacted by kinetic properties leading it to differ from the true equilibrium activity. Moreover, it seems reasonable that some kinetic barriers to extraction or oxidation may depend on membrane physical properties that are themselves correlated with cholesterol content, complicating interpretation of this type of measurement. For example, tighter head-group packing, thicker hydrophobic regions, or a reduction in the number or lifetime of defects could impede access of MβCD or enzymes to cholesterol and slow entry of cholesterol into binding sites. Another, related body of work uses the binding of soluble molecules to membranes to infer cholesterol activity (15, 36, 38, 39). Here too we agree that the localization of these peptides or proteins to membranes via cholesterol binding will depend strongly on the activity of cholesterol, but it will also depend on any other interactions that proteins or peptides make in the membrane bound state. Again, it is reasonable to expect that these other interactions might correlate in complicated ways with cholesterol composition. These caveats do not diminish the value of past conclusions but emphasize that these studies do not directly report on chemical activity in the thermodynamic sense even though this same language is used.

## CONCLUSIONS

Here we demonstrate that μ_c_ within mixtures of cholesterol with one or two phospholipids is far from ideal, meaning that it depends on the full composition of the system rather than just the concentration of cholesterol. Since our measurement of μ_c_ is calibrated, we are able to use results to explore consistency with two models with complimentary treatments of the interactions between the saturated lipids and cholesterol. One model treats cholesterol-saturated lipid interactions as a condensed complex while the other model imposes simple pair-wise interactions, and both models are able to capture experimental trends of μ_c_ with moderate fidelity. Notably, the condensed complex model is found to apply only when the binding energy of the complex is on the order of the thermal energy, indicating that complexes, if present, can readily disassociate on experimental time-scales. Models with pair-wise interactions describe μ_c_ while also qualitatively reproducing regions of liquid-liquid phase coexistence reported in past experiments.

While biological membranes are considerably more complex than these model systems, these measurements provide a valuable perspective on membrane biophysics in cells. They add to a growing body of work that emphasizes the fact that the chemical potentials of cholesterol – or indeed any lipid – should be expected to depend strongly on the other components that are present in the membrane. Since the chemical potential of each lipid is a direct determinant of lipid-protein binding interactions, or the equilibrium lipid exchange between different membranes, we expect these processes to be highly dependent on membrane composition overall.

## AUTHOR CONTRIBUTIONS

SLV devised the central questions of the study. TRS, KW, AG, and SLV developed the experimental protocols. KW and ADG carried out all measurements. SLV analyzed the experimental data in consultation with TRS. TAS and TRS constructed the statistical mechanical models and performed fits to the experimental data in consultation with BM and SLV. SLV and TRS wrote the manuscript with assistance from KW, ADG, TAS, and BM.

## ACKNOWLEDGEMENTS

This study is dedicated to Klaus Gawrisch, whose body of work and general rigorous approach to science set the stage for the current study. SLV thanks Klaus for his longstanding mentorship, his unwavering support, and his regular reminders to stay focused on the big questions.

We thank Artem Ayuyan and Fred Cohen for valuable discussions on the experimental techniques for measuring μ_c_, and Fiona Gaffney for early experimental contributions to this project. This study was supported by grants from the NIH (GM134949 to SLV and GM138341 to BBM) and the NSF (MCB1552439 to SLV).

