## Supplemental Material for "Chemical potential measurements constrain models of cholesterol-phosphatidylcholine interactions"

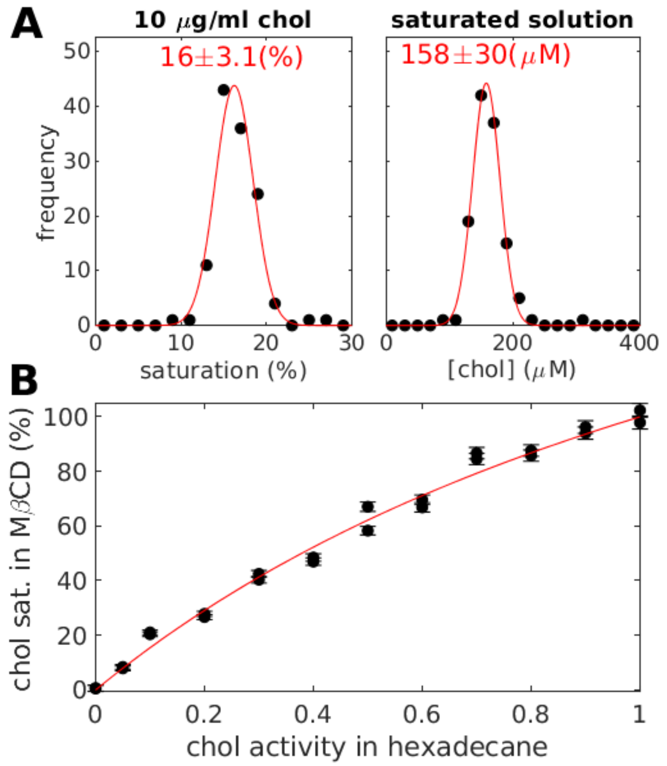


**Supplemental Figure S1: Calibration of saturated solutions of 5mg/ml MβCD and cholesterol.** (A, left) Histogram indicating measurements of the saturation of a 10μg/ml cholesterol standard solution over 122 individual measurements. Saturation is measured against a standard curve with dilutions of a saturated MβCD/cholesterol solution prepared through incubation of 5mg/ml MβCD with cholesterol crystals. (right) Histogram showing the same measurements presented as cholesterol concentration of the saturated solution. Both histograms are fit to a Gaussian distribution to extract the mean and standard deviation shown. (B). Calibration curve relating cholesterol saturation in 5mg/ml MβCD to cholesterol activity. Values are obtained by measuring the saturation of 5mg/ml MβCD solutions equilibrated with hexadecane solutions of known cholesterol activity as described in Methods. Raw values are normalized by 0.93 to account for the minor fraction of hexadecane that interferes with cholesterol binding to MβCD at saturation. The red line is a fit to Eqn. 1.

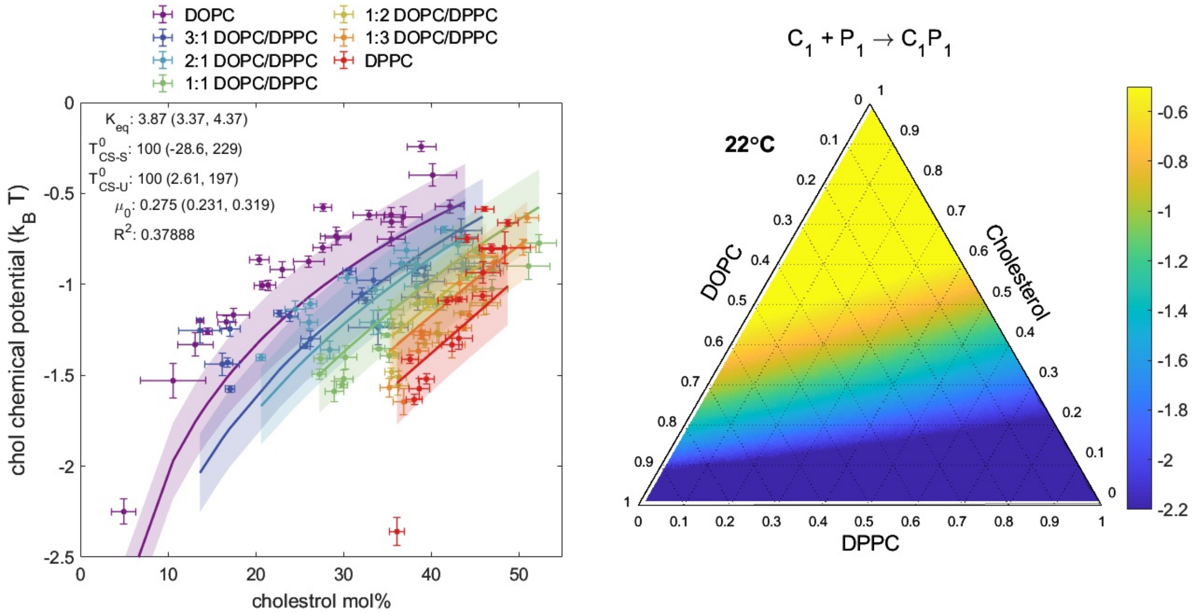


**Supplemental Figure S2: Fitting $\mu_C$ to a regular solutions model with complexes of 1 cholesterol with 1 DPPC molecule, and two repulsive interactions.** This fitted model is identical to the model of Figure 4C, except that an additional repulsion is considered, in this case between complexes and DPPC. The strength of this interaction is given in terms of the critical temperature of a binary mixture of these components: $T^0_{CS-S}$. (left) Experimental data, fitted curves, and shaded confidence intervals as determined by MATLAB's predint. Fitted model parameters, confidence intervals, and Pearson's $R^2$ for this fit are also indicated. (right) Predicted $\mu_C$ surface for all ternary mixtures of cholesterol, DOPC and DPPC, using the fitted model.

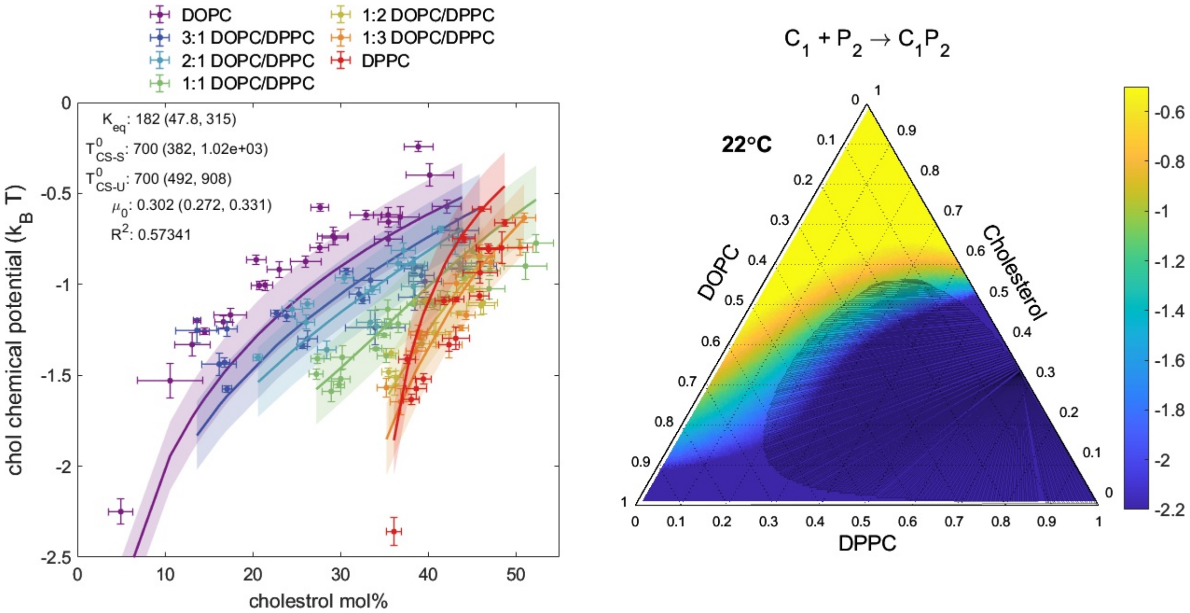


**Supplemental Figure S3: Fitting $\mu_C$ to a regular solutions model with complexes of 1 cholesterol and 2 DPPC molecules, with two repulsive interactions.** Unlike the models of Figure 4, this fitted model uses a 1:2 stoichiometry for the complex CS. An additional repulsion is also considered, in this case between complexes and DPPC. The strength of this interaction is given in terms of the critical temperature of a binary mixture of these components: $T^0_{CS-S}$. (left) Experimental data, fitted curves, and shaded confidence intervals as determined by MATLAB's predint. Fitted model parameters, confidence intervals, and Pearson's $R^2$ for this fit are also indicated. (right) Predicted $\mu_C$ surface for all ternary mixtures of cholesterol, DOPC and DPPC, using the fitted model. The shaded region depicts the approximate liquid-liquid coexistence region of this model.

**Supplementary Table 1: Fitted parameter values and confidence intervals.**

Parameter values and confidence intervals for the five fitted models from Figures 4 and 5. Confidence intervals are for nonlinear least squares fits to the models as described in Methods. MF indicates a mean-field approximation. The interaction parameters for the lattice simulation were obtained by semi-manual optimization, so we did not obtain confidence intervals for those parameters. In addition, each model implies a region for which $\mu_C > 0$, implying spontaneous formation of cholesterol crystals at that such a condition. We consider this to imply a solubility limit for cholesterol, also shown in the table for each model. When they differ, separate values are shown for the Cholesterol-DOPC edge and the Cholesterol-DPPC edge.

| Model: | Shown in Fig: | Fitted parameters (95% confidence interval) | Pearson's $R^2$ | Implied solubility limits of cholesterol (mol %) |
|---|---|---|---|---|
| Ideal solution | 4A | $\mu_C^0$: −0.10 (−0.16, −0.06) $k_BT$ | 0.18 | 110% |
| Ideal solution with complexes | 4B | $K_{eq}$: 5.2 (3.9, 6.5)<br>$\mu_C^0$: 0.36 (0.30, 0.42) $k_BT$ | 0.46 | DOPC edge: 69%<br>DPPC edge: 75% |
| MF Interacting solution with complexes | 4C | $K_{eq}$: 7.2 (5.4, 8.9)<br>$\mu_C^0$: 0.37 (0.31, 0.43) $k_BT$<br>$T_{CS-U}$: 280 (130, 440)K | 0.55 | DOPC edge: 70%<br>DPPC edge: 75% |
| MF Interacting solution without complexes | 5A | $J_{CS}$: − 0.23 (−0.33, −0.13) $k_BT$<br>$J_{CU}$: 0.24 (0.19, 0.29) $k_BT$<br>$J_{SU}$: 0.59 (0.44, 0.76) $k_BT$<br>$\mu_C^0$: 0.06 (−0.04, 0.16) $k_BT$ | 0.78 | DOPC edge: 91%<br>DPPC edge: 95% |
| Lattice simulation of Interacting solution without complexes | 5B | $J_{CS}$: − 0.62 $k_BT$<br>$J_{CU}$: 0 $k_BT$<br>$J_{SU}$: 1.06 $k_BT$<br>$\mu_C^0$: 0.54 $k_BT$ | 0.50 | DOPC edge: 53%<br>DPPC edge: 68% |

**Supplementary note 1: computation of chemical potentials in a mole fraction basis**

While chemical potentials are typically defined in terms of partial derivatives with respect to number of each component, it is convenient here to compute derivatives with respect to mole fractions instead. This supplementary note derives expressions for the chemical potentials in terms of those derivatives.

For a system containing $r$ molecular species, standard definitions of Gibbs free energy $G$ and the chemical potentials give (assuming const. pressure and temperature for simplicity):

$$dG(N_1, N_2, \dots, N_r) = \sum_{i=1}^{r} \mu_i dN_i.$$

It is convenient to rewrite this in terms of $N = \sum_i N_i$, and $x_i = N_i/N$, for $i = 2, \dots, r$. For consistency, $x_1 = 1 - \sum_{i>1} x_i$ is taken to be dependent on the other $x_i$, and is no longer a formal variable of $G$. Note that the choice of the first component as the dependent variable is arbitrary, and the following forms apply whichever component is chosen. However, it is important to keep track of which one is dependent, because the partial derivatives obtained below are dependent on this choice. Then we have

$$dN_i = Ndx_i + x_i dN, i > 1$$

$$dN_1 = dN - \sum_{i>1} dN_i$$

$$= dN\left(1 - \sum_{i>1} x_i\right) - N\left(\sum_{i>1} dx_i\right).$$

Rewriting the exact differential in terms of the new variables, we obtain:

$$dG(N, x_2, \dots, x_r) = \sum_i \mu_i x_i \, dN + N \sum_{i>1} (\mu_i - \mu_1) \, dx_i$$

We can then read off the partial derivatives with respect to the new variables. The derivatives with respect to $x_i, i > 1$ now give differences of chemical potentials:

$$\frac{\partial(G/N)}{\partial x_i} = \mu_i - \mu_1, \;\; i > 1$$

The final chemical potential can be obtained from the fact that $G$ is an extensive function, so is proportional to $N$ (at fixed mole fractions of the components):

$$\frac{\partial G}{\partial N} = \sum_i \mu_i x_i = \frac{G}{N}.$$

Putting this together with the above and defining $g = G/N$, we may conclude:

$$g - \sum_{i>1} x_i \frac{\partial g}{\partial x_i} = \mu_U \left(1 - \sum_{i>1} x_i\right) + \mu_U \sum_{i>1} x_i = \mu_1.$$

This expression for $\mu_1$ can then be substituted into the above difference to obtain expressions for the $\mu_i, i > 1$:

$$\mu_i(x_2,\dots,x_r) = g(x_2,\dots,x_r) + \frac{\partial g}{\partial x_i} - \sum_{i>1} x_i \frac{\partial g}{\partial x_i}, i > 1.$$

These expressions will be used in the following to derive the function forms of the relevant chemical potentials of specific models.

**Supplementary note 2: Regular solution models**

The theoretical models discussed in the main text take the form of regular solution models. A regular solution model for a mixture considers nearest-neighbor interactions and internal degrees of freedom of each of a set of components, while neglecting oriented interactions. Define the Hamiltonian for such a model by

$$\mathcal{H} = \sum_{i=1}^{r} \mu_i^0 + \sum_{i<j} N_{ij} J_{ij},$$

where $N_i$ indicates the number of species $i$, $\mu_i^0$ is the free energy per molecule of the internal degrees of freedom for species $i$, $N_{ij}$ is the number of $i-j$ nearest-neighbor pairs, and $J_{ij}$ specifies the interaction energy of such a pair. Note that we need not include $i-i$ interactions explicitly, because by an appropriate transformation of the $J_{ij}$ and $\mu_i^0$, the $J_{ii}$ can be set to 0 without altering the Hamiltonian overall(1).

In a mean field, sometimes known as zeroth order, approximation, we take the entropy of mixing to be that of an ideal mixture, and the average energy to be the energy of a well-mixed configuration, so that the free energy per molecule becomes

$$g_N = \frac{G}{N} = \sum_i \{k_B T\, x_i \log x_i + x_i \mu_i^0\} + z \sum_{i<j} x_i x_j J_{ij},$$

where, as above, $x_i = N_i/N$ is the mole fraction of each component, and $z$ is the coordination number of the lattice in question, which we take to be 4 in all of our models. In some cases, such as the models that include cholesterol-phospholipid complexes that are described in Supplementary Note 4, where all non-zero interactions are repulsive, we find it convenient to rewrite the interaction parameters as (mean-field) critical temperatures for phase separation, by the transformation $zJ_{ij} = 2k_B T_{i-j}$, where $T_{i-j}$ is then the critical temperature for phase separation of binary $i-j$ mixtures.

**Supplementary note 3: Mean field regular solution without complexes**

The model that is fitted in Figure 5A corresponds to a regular solution, defined as above, with three components only, C, S, and U, corresponding to cholesterol, DPPC (saturated phospholipid), and DOPC (unsaturated phospholipid). All of the interaction parameters are allowed to vary freely: $J_{CS}$, $J_{CU}$ and $J_{SU}$. We consider the regular solution free energy as a function of $x_S$ and $x_U$ and use the chemical potential identities derived above to obtain:

$$\frac{\mu_C}{k_B T} = \frac{1}{k_B T}\left( g - x_S \frac{\partial g}{\partial x_S} - x_U \frac{\partial g}{\partial x_U} \right)$$

$$= \log x_C + \frac{\mu_C^0}{k_B T} + z \frac{J_{CS}}{k_B T}(x_S^2 + x_u x_S) + z \frac{J_{CU}}{k_B T}(x_U^2 + x_U x_S) - z \frac{J_{SU}}{k_B T}(x_S x_U)$$

**Supplementary note 4: mean field regular solution models with complexes**

Following McConnell and Radhakrishnan(2–4), we model a lipid bilayer composed of cholesterol, a saturated lipid (DPPC), and an unsaturated lipid (DOPC). As above, C, S, and U are used as shorthands for cholesterol, DPPC (saturated lipid) and DOPC (unsaturated lipid), respectively. A condensed complex written CS is allowed to form between cholesterol and DPPC in a reversible reaction assumed to have $q\colon p$ stoichiometry with equilibrium constant $K_{eq}$.

$$qC + pS \overset{K_{eq}}{\longleftrightarrow} C_q S_p$$

For later convenience we distinguish $N_i$, the total number of lipid species $i \in \{C, S, U\}$ in the system before the reaction, from $N_i{}'$, the number of species $i$ after the reaction comes to equilibrium. Furthermore, $N = N_C + N_S + N_U$, and $x_i = N_i/N$, and $N' = N_C' + N_S' + N_U' + N_{CS}$, and $x_i' = N_i'/N'$. As above, we begin with the regular solution free energy per molecule ($N'$) of an equilibrium mixture of C, S, U, and CS:

$$g_{N'}^{RS} = \frac{G}{N'} = \sum_i x_i'(k_B T \ln x_i') + 2k_B \sum_{i<j} x_i' x_j' T_{i-j}^0$$

Where $\mu_i^0$ is standard chemical potential of pure component $i$, $x_i'$ is equilibrium mole fraction of $i$ , and $T_{ij}^0$ is the critical temperature of the $i - j$ binary pair.

To account for the complexation reaction we add to this a term:

$$G_{complex} = N_{CS} \Delta G_{complex} = -k_B T N_{CS} \ln K_{eq},$$

and finally allow for a non-zero $\mu_C^0$ in order to compare to refer the chemical potential to the energy scale set by taking the cholesterol crystal as standard state:

$$G_{standard} = N_C' \mu_C^0$$

Finally, following (4), we assume all critical temperatures are zero except $T_{cs-s}^0$ (immiscibility between CS and S) and $T_{cs-u}^0$ (immiscibility between CS and U).

Following these claims, the free energy per molecule (of which there are $N'$ in number) denoted as $g'$ takes the form

$$\frac{g'}{k_B T} = \left( x_c' \,(\mu_C^0/\mathrm{k_B T} + \ln x_c') + x_s' \ln x_s' + x_u' \ln x_u' + x_{cs} \ln \frac{x_{cs}}{K_{eq}} \right) + \frac{2x_{cs}}{T} (T_{cs-s} x_s' + T_{cs-u} x_u')$$

(1)

It is useful to be able to write the free energy in terms of the mole fractions that are inserted at the beginning of the experiment $x_i$, rather than the ones after the reaction reaches equilibrium $x_i'$. Begin by defining the system in terms of a reaction progress parameter $\gamma$ which is minimized in all calculations of the total free energy $G^*$

$$\gamma = \frac{N_{cs}}{N_c' + N_s' + N_u' + (q+p)N_{cs}}$$

We can write the transform in terms of this parameter and a parameter $\lambda$ as follows

$$x_{cs} = \gamma\lambda$$

$$x_{c'} = (x_c - q\gamma)\lambda$$

$$x_{s'} = (x_s - p\gamma)\lambda$$

$$x_{u'} = x_u\lambda$$

$\lambda$ can then be determined by noting our constraints: $x_c + x_s + x_u = x_{c'} + x_{s'} + x_{u'} + x_{cs} = 1$. Solving with these constraints yields.

$$\lambda = [1 + (1 - q - p)\gamma]^{-1}$$

Using the same constraints, we can solve for gamma in terms of $x_{cs}$

$$\gamma = \frac{x_{cs}}{1 + (p + q - 1)x_{cs}}$$

And then use these two relations to eliminate $\gamma$ and $\lambda$ from our transformations entirely

$$x_{c'} = x_c[1 + (p + q - 1)x_{cs}] - qx_{cs}$$

$$x_{s'} = x_s[1 + (p + q - 1)x_{cs}] - px_{cs}$$

$$x_{u'} = (1 - x_c - x_s)[1 + (p + q - 1)x_{cs}]$$

Because the number of particles in the system is not conserved through the reaction it is useful to know how the initial number of molecules inserted into the system $N$ relates to the final number $N'$. We can use our relation for $x_u'$ combined with the fact that $N_u' = N_u$ to find this relation.

$$x_u' = x_u\lambda$$

$$x_U{}' = x_U\lambda$$

$$\lambda = \frac{N}{N'}$$

It is worth underlining that with this relation, $g = G/N$ may be obtained from $g' = G/N'$ by

$$g = g'\lambda^{-1}$$

*Letting the reaction equilibrate*:

To extract information from $g$ or $g'$ we need to minimize $G$ with respect to the fraction of complex $x_{cs}$. This is equivalent to minimizing $g = G/N$, because $N$ does not vary as the reaction progresses. So we define $g^* = g(x_c, x_s, x_{cs}^*)$ such that

$$\left.\frac{\partial g}{\partial x_{cs}}\right|_{x_{cs}=x_{cs}^*} = \lambda^{-1}\left.\frac{\partial g'}{\partial x_{cs}}\right|_{x_{cs}=x_{cs}^*} = 0$$

Unfortunately computing such an $x_{cs}^*$ leads to a transcendental equation which does not have an analytical solution. Thus, we compute $x_{cs}^*$ numerically using MATLAB's numerical minimization routine fminbnd.

*Calculating $\mu_c'$:*

With numerical $x_{cs}^*$, we may return to general stoichiometry $q\!:\!p$ and the primed basis for simplicity to calculate the chemical potential of free cholesterol (now ignoring the denominator $k_B T$) using the relations of Supplementary Note 1:

$$\mu_{c'} - \mu_{u'} = \left(\frac{\partial g^*}{\partial x_c'}\right)_{T,P,x_{i\neq c}'}$$

$$\mu_{u'} = g^* - \sum_{i\neq u} x_i' \frac{\partial g^*}{\partial x_i'}$$

We can rewrite our expression for ${\mu_c}'$ as

$$\mu_{c'} = \left(\frac{\partial g^*}{\partial x_c'}\right)_{T,P,x_{i\neq c}'} + \mu_{u'}$$

$$\mu_{c'} = \left(\frac{\partial g^*}{\partial x_c'}\right)_{T,P,x_{i\neq c}'} + g^* - x_s' \frac{\partial g^*}{\partial x_s'} - x_c' \frac{\partial g^*}{\partial x_c'} - x_{cs}^* \frac{\partial g^*}{\partial x_{cs}^*}$$

$$\mu_{c'} = g^* + (1 - x_c') \frac{\partial g^*}{\partial x_c'} - x_s' \frac{\partial g^*}{\partial x_s'} - x_{cs}^* \frac{\partial g^*}{\partial x_{cs}^*}$$

Where $g^*(x_c', x_s', x_{cs}^*)$ in units of $k_B T$ is

$$g^* = \left( x_c'(\mu_C^0 + \ln x_c') + x_s' \ln x_s' + (1 - \mathrm{x}_\mathrm{c}' - x_s' - x_{cs}^*) \ln(1 - \mathrm{x}_\mathrm{c}' - x_s' - x_{cs}^*) + x_{cs}^* \ln \frac{x_{cs}^*}{K_{eq}} \right) + \frac{2x_{cs}^*}{T} \left( T_{cs-s} x_s' + T_{cs-u}(1 - \mathrm{x}_\mathrm{c}' - x_s' - x_{cs}^*) \right)$$

Proceeding by first looking at each partial derivative in $\mu_{c'}$

$$\frac{\partial g^*}{\partial x_c'} = \mu_C^0 + \log\left(\frac{x_c'}{1 - x_c' - x_s' - x_{cs}^*}\right) - \frac{2x_{cs}^*}{T} T_{cs-u}$$

$$\frac{\partial g^*}{\partial x_s'} = \log\left(\frac{x_s'}{1 - x_c' - x_s' - x_{cs}^*}\right) + \frac{2x_{cs}^*}{T} (T_{cs-s} - T_{cs-u})$$

$$\frac{\partial g^*}{\partial x_{cs}^*} = \log\left(\frac{x_{cs}^*}{K_{eq}(1 - x_c' - x_s' - x_{cs})}\right) + \frac{2x_s'}{T} T_{cs-s} - \frac{2(x_c' + x_s' + 2x_{cs}^* - 1)}{T} T_{cs-u}$$

Then combining these results, we arrive at

$$\mu_{c'} = \mu_\mathrm{C}^0 + \log(x_c') - \frac{2x_{cs}^*}{T} [(1 - x_c' - x_s' - x_{cs}^*) T_{cs-u} + x_s' T_{cs-s}]$$

Note that for boundary data (where $x_s$ or $x_u = 0$ ) we need to define separate free energies $g^*_{\mathrm{DPPC}}$ and $g^*_{\mathrm{DOPC}}$ in the limit as the relevant mole fraction goes to 0, yielding ultimately:

$$\mu_{c'}^{\mathrm{DPPC}} = \mu_C^0 + \log(x_c') - \frac{2x_{cs}^* x_s'}{T} T_{cs-s}$$

$$\mu_{c'}^{\mathrm{DOPC}} = \mu_C^0 + \log(x_c')$$

In equilibrium $\mu_{c'}$ will be equivalent to $\mu_c$ giving us a final piece-wise expression for $\mu_c$ as a function of the $x_s$, $x_c$, $x_{cs}$ basis (but written here in the $x_i'$ basis for brevity):

$$\mu_c = \begin{cases} \mu_C^0 + \log(x_c') & x_s = 0 \\ \mu_C^0 + \log(x_c') - \frac{2x_{cs}^*}{T}[(1 - x_c' - x_s' - x_{cs}^*)T_{cs-u} + x_s' T_{cs-s}] & x_i \neq 0 \\ \mu_C^0 + \log(x_c') - \frac{2x_{cs}^* x_s'}{T} T_{cs-s} & x_u = 0 \end{cases}$$

**Supplementary note 5: Estimating chemical potentials from lattice model simulations of the regular solution model without complexes**

The model that is shown in Figure 5B corresponds to the same model as Figure 5A, but without making the mean field approximation. Therefore, we cannot write an explicit expression for the free energy or its derivatives, and have to estimate these from the simulations themselves. Simulations are carried out as described in Methods, and then analyzed as follows.

Estimation of derivatives $\partial g \, / \, \partial x_i$ in a lattice model using a Monte Carlo scheme:

Consider a Monte Carlo update scheme that allows components from the lattice simulation to be exchanged with a bath of particles. It must satisfy detailed balance in order for it to equilibrate. This means

$$P_{A\to B} P_A = P_{B\to A} P_B$$

Where *A*, *B* indicate states of the system, $P_A$ is the probability of state *A* in equilibrium, and $P_{A\to B}$ is the probability under the Monte Carlo scheme of a state transition from *A* to *B*. The equilibrium distribution is given by the grand canonical ensemble

$$P_A \propto \exp\left(-\beta\left[E_A - \sum_k \mu_k N_k{}^{(A)}\right]\right)$$

Where $E_A$ indicates the (total) energy of state $A$, $k$ indexes the chemical species of the mixture, $\mu_k$ is the chemical potential of species $k$, and $N_k^{(A)}$ is the number of molecules of species $k$ that are present in state *A*. The proportionality constant is the grand partition function, which need not be estimated here because we will only need ratios of probabilities.

We will only consider schemes with moves that swap the species that is present at a random single site $i$. Write $(s_i, k \to l)$ to denote such a swap. For such a swap, the ratio $P_B/P_A$ only depends on the interaction energies between site $i$ and its four neighbors. In particular,

$$\frac{P_{s_i,l\to k}}{P_{s_i,k\to l}} = \frac{P_{s_i=k}}{P_{s_i=l}} = \exp(-\beta[(E_k - E_l) - (\mu_k - \mu_l)]),$$

where the first equality comes from rearranging the detailed balance condition, and the second comes from the grand canonical ensemble. $E_k$ now indicates the sum of the interactions of site $i$ with its neighbors if the species at that site is $k$. The energy of the rest of the lattice need not be considered, because it does not depend on the species at site $i$, and thus cancels in the ratio.

One proposal for transition probabilities (conditioned on site $i$ being chosen as the random site to swap) that satisfies the above condition is

$$P_{s_i,k\to l} = C \cdot \exp(-\beta[(E_l - E_k) - (\mu_l - \mu_k)]/2)$$

with the constant $C$ chosen so that no transition probability exceeds 1. Then

$$\frac{P_{s_i,l\to k}}{P_{s_i,k\to l}} = \frac{\exp(-\beta[(E_k - E_l) - (\mu_k - \mu_l)]/2)}{\exp(-\beta[(E_l - E_k) - (\mu_l - \mu_k)]/2)} = \exp(-\beta[(E_k - E_l) - (\mu_k - \mu_l)])$$

as required. Consider the simulations as a sample from the equilibrium distribution, i.e. under the assumption that the $\mu_k$have been chosen such that the average number of molecules of species $k$, $\langle N_k \rangle$ under the grand canonical ensemble is equal to the (fixed) number that are actually present in the given simulation. Consider the sum

$$\begin{aligned} W_{k\to l} &= \sum_i \delta(s_i = k)\exp(-\beta[E_l - E_k]/2) \\ &= \exp(-\beta(\mu_l - \mu_k)/2)\sum_i \delta(s_i = k)\exp[-\beta[(E_l - E_k) - (\mu_l - \mu_k)]/2] \\ &\propto \exp(-\beta(\mu_l - \mu_k)/2) P_{s_i,k\to l} P_{s_i=k} \end{aligned}$$

Therefore, by taking advantage of detailed balance, we can conclude that

$$\frac{W_{k\to l}}{W_{l\to k}} = \exp\big(\beta(\mu_k - \mu_l)\big)\frac{P_{s_i,k\to l}P_{s_i=k}}{P_{s_i,l\to k}P_{s_i=l}} = \exp\big(\beta(\mu_k - \mu_l)\big).$$

so that

$$\mu_k - \mu_l = k_B T \log\left(\frac{W_{k\to l}}{W_{l\to k}}\right).$$

To obtain the chemical potentials themselves from these differences, note that supplementary note 1 gives us a way to write these differences as partial derivatives of the free energy per molecule $g$. For the three species C,S and U, taking as dependent mole fraction $x_U = 1 - x_C - x_S$, we have

$$\frac{\partial g(x_C, x_S)}{\partial x_C} = \mu_C - \mu_S.$$

Then we can estimate the overall free energy by numerically integrating these partial derivatives:

$$g(x_C, x_S) = \mu_0 + \int_{x_C^*}^{x_C} \frac{\partial g}{\partial x_C}(x, x_S^*)dx + \int_{x_S^*}^{x_S} \frac{\partial g}{\partial x_S}(x_C, x)dx.$$

where $x_C^*$ and $x_S^*$ are chosen as the origin of the integration and $\mu_0$ is an integration constant. We evaluate the derivatives using simulations that are closely spaced in composition space, with spacing of at most .05 in both $x_C$ and $x_S$. At the edges of composition space where one component vanishes, finer spacing is chosen. These samples are then interpolated using cubic spline interpolation to obtain the derivatives at arbitrary interior points of composition space. Integration is carried out as a Riemann sum with spacing of at most .002 in $x_C$ and $x_S$.

Finally, with $g$ and the various $\mu_i - \mu_j$ in hand, we can compute for any composition:

$$\mu_C = g - x_U(\mu_U - \mu_C) - x_S(\mu_S - \mu_C)$$

as derived in Supplementary Note 1.
